# Multi-Omics Characterization of ctDNA Release Mechanisms in Ovarian Cancer

**DOI:** 10.1101/2025.08.03.668326

**Authors:** Mai T.N. Nguyen, Yilin Li, Daria Afenteva, Kari Lavikka, Giovanni Marchi, Johanna Hynninen, Sampsa Hautaniemi, Jaana Oikkonen

**Affiliations:** Research Program in Systems Oncology, Faculty of Medicine, University of Helsinki, Helsinki 00291, Finland; Department of Obstetrics and Gynecology, Turku University Hospital and University of Turku, Kiinamyllynkatu 4, Turku 20521, Finland

**Keywords:** ctDNA release biology, multi-omics, tumor intrinsic mechanisms, ovarian cancer, HGSC

## Abstract

Circulating tumor DNA (ctDNA) is a powerful biomarker capable to predict tumor dynamics and treatment response. Despite its importance, the biological mechanisms behind ctDNA release have remained unclear. The variability among patients is affected by cancer burden, histology and stage, but these factors cover only a part of the detected variability. Herein, we characterized molecular drivers of baseline ctDNA variability in a real-world cohort of 118 patients with ovarian high-grade serous carcinoma (HGSC). Genomic and transcriptomic analyses revealed a strong positive correlation between ctDNA levels and cellular proliferation, and an inverse relationship with immune activity. Particularly, low ctDNA tumors exhibited higher bulk expression of *mucins* and *CIITA*, suggesting that intrinsic immune responses and immune landscape are linked to ctDNA release. The low and high ctDNA levels were significantly associated with poor prognosis compared to medium level in unresectable HGSC patients treated with neoadjuvant chemotherapy. In summary, our results suggest that ctDNA release is modulated by cancer cell proliferation and tumor microenvironment.

## Introduction

High-grade serous carcinoma (HGSC) is the most abundant and lethal epithelial ovarian cancer with 5-years survival of less than 40% ^1^. The use of ctDNA in treatment response monitoring and guidance in HGSC have been shown in many studies ^2–6^. However, the utility of ctDNA is greatly influenced by the amount of tumor DNA fragments in the bloodstream ^7–9^. Although HGSC is generally diagnosed at an advanced stage, with extensive intra-abdominal spread and a high tumor burden, the amount of ctDNA is typically low. For example, undetectable ctDNA has been reported in 36% of patients ^10^. A deeper understanding of the biological mechanisms influencing ctDNA release is crucial to improving its reliability as a clinical biomarker.

The release of ctDNA is believed to occur primarily through cancer cell death processes, such as apoptosis, necrosis, and autophagy ^11–13^, as well as through active mechanisms like exosome secretion ^14^. Previous studies have suggested a correlation between ctDNA levels and tumor burden, with higher levels observed in more advanced cancers ^14,15^. Additionally, ctDNA abundance varies across cancer types ^16^, reflecting tumor-specific mechanisms affecting either its release from tumors or clearance from circulation ^14,17,18^. Furthermore, ctDNA can be concealed by the abundance of cell-free DNA (cfDNA) from other sources than tumors, mainly immune cells ^19^. Studies in lung ^20^, colorectal ^21^, and breast ^22^ cancers have indicated that tumors with higher proliferation rate tend to exhibit increased ctDNA release, although these analyses often rely on binary classifications of ctDNA positivity rather than quantitative assessments.

Despite these findings, a significant portion of individual variability in ctDNA abundance remains unexplained even at advanced stage cancers such as HGSC, which poses challenges for the clinical utility of ctDNA, particularly in the areas of personalized medicine and cancer detection ^7,10,23^. This limitation is especially important given the increasing reliance on ctDNA positivity as a selection criterion in many clinical trials ^23–25^. Therefore, a better understanding of ctDNA release, particularly in patients with very low ctDNA fractions, is essential for improving its application in oncology.

To address this gap in HGSC patients, we leveraged ctDNA, whole-genome sequencing (WGS) and RNA-seq data from 118 patients with HGSC who belong to the longitudinal, multi-region, observational, prospective DECIDER trial ^26^. Our results, based on one of the largest real-world multi-modal data in ovarian cancer to date, show that ctDNA release in HGSC is governed by biological differences in tumors, a complex interplay of proliferation, metabolic activities, and immune responses.

## Materials and Methods

### Patient recruitment

We collected plasma samples from 128 patients (Supplementary Table S1) recruited at Turku University Hospital, Finland, between 2015 and 2022. All patients belong to the DECIDER cohort (Multi-Layer Data to Improve Diagnosis, Predict Therapy Resistance and Suggest Targeted Therapies in high-grade serous carcinoma (HGSOC); ClinicalTrials.gov identifier <u>NCT04846933</u>). Among them, 10 patients were diagnosed with benign ovarian lesions, and 118 were diagnosed with HGSC (Supplementary Table S1). The wellbeing services county of Southwest Finland ethics committee approved the study (VARHA/28314/13.02.02/2023), and informed consent was obtained from all patients. Patient clinical data were collected from electronic hospital records (EHR), manually filled structured surgical operation forms, and by interviewing the patients.

### Plasma sample collection

Pretreatment blood samples (10ml) were collected into EDTA tubes from all patients and centrifuged twice for 10 minutes at 2000xg. Centrifuged plasma was stored at −80°C in 1ml subsamples within 2 hours after collection for cell free DNA (cfDNA) extraction. CfDNA was extracted using the QIAmp Circulating Nucleic Acid Kit (Qiagen, #55114), and its concentration was measured with a Qubit fluorometer (Thermo Fisher Scientific) and Agilent 5400 fragment analyzer system (Agilent Technologies). Library preparation was done with NEBNext Ultra II DNA Library Prep Kit skipping the fragmentation step (New England Biolabs, #E7370L). Libraries were purified with AMPure XP and sequenced on the Illumina NovaSeq 6000 system (150bp paired end sequencing, Illumina Inc.). Library preparation and sequencing was done at Novogene Company Ltd (UK). The median DNA amount extracted for sequencing was 51ng (18-63). The ctDNA samples underwent shallow whole-genome sequencing (sWGS) with a mean coverage of 0.4 (0.25–0.90).

Additionally, whole exome sequencing (WES) was performed for part of the samples with Agilent SureSelect Human All Exon V6, 600x coverage. Part of these samples were sequenced at Biomarker Technologies BMK GmbH (Germany) where extraction was performed with Quick-cfDNA™ Serum & Plasma Kit (Zymo Research). Library preparation used the same NEBNext Ultra II DNA library prep kit as sWGS protocol.

Plasma samples had been collected between 2016 and 2022, there were no significant differences in DNA amount extracted and ctDNA fraction estimated across that period of collecting time (Supplementary Figure S1).

### Whole blood control and tumor tissue samples

Whole blood control and tumor tissue samples were obtained and processed as described in our previous study ^26^. Briefly, blood control samples were collected at the initiation of treatment, and DNA extraction was performed at the Auria biobank. Tumor tissue samples were obtained during laparoscopy and debulking surgery, with simultaneous extraction of DNA and RNA using the Qiagen AllPrep kit (Qiagen, #80204). Subsequently, samples were sent for library preparation and sequencing. Samples were sequenced with either Illumina NovaSeq 6000 (Novogene (UK) Company Ltd), Illumina HiSeq X Ten, MGISEQ 2000 or BGISEQ-500 (BGI Europe A/S, Denmark) employing paired end sequencing. The whole-genome sequencing (WGS) and RNA sequencing (RNA-seq) data was available for 100 (Supplementary Table S2) and 102 patients (Supplementary Table S3), respectively.

### Preprocessing sWGS, WGS, WES, and RNA-seq data

Preprocessing of WGS, sWGS, WES and RNA-seq data, including quality control (FastQC), trimming (Trimmomatic), and alignment, are described in detail in previous studies ^26,27^, and available through https://zenodo.org/records/7852210. Expression gene level counts were calculated after the alignment.

### ctDNA fraction estimation with ichorCNA for sWGS data

Copy-number profiles were used to estimate ctDNA fraction for all sWGS samples with ichorCNA ^28^. Copy-numbers were estimated from 1Mb read count windows. GATK ^29^ was used to preprocess these windows by PreprocessingIntervals after which GC content and mappability were annotated by AnnotateInvervals. Read counts were calculated by function calculateBamCoverageByInterval of R package PureCN (version 2.2.0) ^30^. Regions with mappability smaller than 0.9 were excluded. Parameter tuning in ichorCNA followed the author’s recommendation of low tumor fraction (https://github.com/broadinstitute/ichorCNA/wiki/Parameter-tuning-and-settings). To construct panel of normal for ichorCNA, we downsampled the white blood control samples to a lower coverage by fraction of 0.01 with samtools ^31^. The pipeline was run with Anduril ^32^.

To evaluate the reliability of ctDNA fraction estimated by ichorCNA ^28^, we used manually curated truncal *TP53* variants from WGS tissue data and estimated their variant allele frequency (VAF) from matched ctDNA WES as control values (Supplementary Method). Almost all HGSC patients have truncal *TP53* mutation, and it has been used as a biomarker for tumor fraction in HGSC ^33^. With our pipeline, the estimated ctDNA fraction values correlated very well with the *TP53* VAF (Supplementary Figures S2, S3). Especially for low ctDNA fraction samples, our panel of normals (PoN) produced higher precision with coefficient of 1 and 0.92 (Supplementary Figures S2b, S3b) for samples with *TP53* VAF of smaller than 0.03 and 0.065, respectively, higher than the original ichorCNA PoN with the coefficient of 0.47 and 0.65 (Supplementary Figures S2a, S3a), which allowed us to estimate ctDNA level in larger fraction of samples.

### Mutation calling and signatures

Somatic short variant and copy-number calling for tumor tissue samples were described in our previous study ^26^. Briefly, somatic short variants were called with Mutect2 ^34^. Copy-numbers were called with GATK ^29^, and purity and ploidy were estimated with modified ASCAT ^26,35^. WGS data from 191 solid tumor tissue samples were available for 100 patients. Purity of included WGS samples were in a range of 0.08–1.00 (Supplementary Table S2).

Mutational signatures were fitted for pretreatment tissue samples utilizing COSMIC ^36^ v3.3.1 reference signatures (https://cancer.sanger.ac.uk/signatures/downloads/). The signatures were adjusted for GRCh38 nucleotide frequencies, excluding the Y and mitochondrial chromosomes. There were three types of mutations in this analysis including: single base substitutions (SBS), double base substitutions (DBS), and insertions or deletions (ID). For group comparisons, we opted for the highest purity sample per patient. For survival analyses, all samples were used. Specifically for the signatures SBS3 and ID5, we classified the values into either zero or non-zero categories.

In the mutational burden analysis and mutational signatures, we excluded variants identified in all samples and tagged as artifacts by the GATK FilterAlignmentArtifacts model. For copy-number variant analysis, we calculated the average number of breakpoints per chromosome. This was done by dividing the total number of breakpoints identified in each sample by the genome size.

Additionally, homologous recombination status defined in our previous study ^37^ was categorized as homologous recombination deficiency (HRD), proficiency (HRP).

### Transcriptomics data from bulk RNA sequencing

Preprocessing and decomposition for bulk RNA samples were described in our previous study ^27^. Briefly, RNA data were decomposed with PRISM ^27^ into cell-type-specific whole-transcriptome profiles. Three main components in this analysis included epithelial ovarian cancer (EOC), immune, and fibroblast cells. For our downstream analysis, we used transcriptomic profiles of the EOC component that were available for 245 samples from 102 patients (Supplementary Table S3).

To validate the reliability of the cancer cell component decomposition from bulk RNA via PRISM, we compared the EOC proportion from bulk RNA sequencing of solid tissue samples with purity estimates from copy-number calling from matched WGS samples. The high correlation (*R* = 0.83, *p*-value < 1*10^-5^) (Supplementary Figure S4) between these tumor content estimates from 97 patients ensured the robustness of bulk RNA decomposition.

### RNA-seq differential expression and pathway enrichment analysis

Initially, gene set enrichment analysis (GSEA) was performed using the R package GSVA ^38^ (v1.52.3) to calculate pathway scores at the sample level, with a minimum gene set size of 20. The pathway scores for each patient were determined as the mean of scores from all samples per patient. The Hallmark pathways from the Molecular Signatures Database (MsigDB) ^39^ were employed for the pathway analysis.

The differential expression gene (DEG) analysis between two groups was conducted using DESeq2 ^40^ (v1.44.0) on samples with the highest EOC proportion from each patient. Subsequently, GSEA of the DEG results was performed using fgsea (https://github.com/ctlab/fgsea/, v1.30.0). This analysis utilized ordered DE t-statistics, involving 10,000 permutations, with the minimum and maximum gene set sizes set to 20 and 500, respectively.

### Kaplan–Meier and Cox regression analysis

For survival analyses using progression-free interval (PFI), we filtered out patients that either received only a single chemo cycle or the outcomes could not be evaluated, resulting in 114 patients (Supplementary Table S1). R packages survival (v3.6.4), survminer (v0.4.9), and finalfit (v1.0.7) were employed for conducting these analyses.

### Multiple groups comparison

For group comparisons we employed the Wilcoxon rank-sum test for comparing two groups and the Kruskal–Wallis test for assessing differences among multiple groups. The proportional differences were studied using χ^2^ test. These statistical tests were chosen to ensure a comprehensive assessment of differences between groups, considering the nature of the data and the potential variability within and between groups.

### Other statistical and visualization analyses

We employed ggplot2 (v3.5.1), ggpubr (v0.6.0), pheatmap (v1.0.12), and corrplot (v0.92) packages for the visualizations. Biorender (https://www.biorender.com) was used for illustrations. The principal-component analysis (PCA) was performed with R package factoextra (v1.0.7). All analyses and visualizations were performed in R 4.4.1.

## Results

### Patients cohort and ctDNA fraction estimation

Our study included samples from 118 patients with HGSC (median age: 71) and 10 benign controls, with all diagnoses confirmed by pathologists. The majority of HGSC patients (88%) were diagnosed at advanced stages (FIGO stage III: *n* = 85, stage IV: *n* = 19, Table 1). Among these patients, 38% received neoadjuvant chemotherapy (NACT, *n* = 45), while 60% underwent primary debulking surgery (PDS, *n* = 72). One patient, who received a single chemotherapy cycle before surgery, was categorized under ‘other treatment.’ Additionally, homologous recombination (HR) status was determined for HGSC patients based on the classification by Koskela et al. ^37^. Of the cohort, 36% were classified as homologous recombination deficient (HRD, *n* = 43), and 57% were homologous recombination proficient (HRP, *n* = 67). HR status was unavailable for 8 patients (Table 1).

**Table 1:** Clinical characteristics of the patient cohort.

| Clinical factors | *N* |
| --- | --- |
| Benign | **10** |
| HGSC | **118** |
| FIGO Stages |  |
| I | 3 |
| II | 11 |
| III | 85 |
| IV | 19 |
| Treatment |  |
| NACT | 45 |
| PDS | 72 |
| Other | 1 |
| Homologous recombination status |  |
| HRD | 43 |
| HRP | 67 |
| NA | 8 |
| Age (years) | 41 - 88 |

HGSC: High-grade serous carcinoma; NACT: Neoadjuvant chemotherapy; HRD: Homologous recombination deficiency; HRP: Homologous recombination proficient; FIGO: International Federation of Gynecology and Obstetrics;

Baseline ctDNA fraction was estimated using ichorCNA, based on CNV profiles obtained from shallow whole-genome sequencing of treatment-naïve plasma samples (*n*=128). To estimate the accuracy of the ctDNA fractions, we performed linear regression between *TP53* mutation-based estimates from high-coverage sequencing and the ichor-CNA-based estimates from the low-coverage sequencing which demonstrated concordance even in samples with low ctDNA fractions (Supplementary Methods, Supplementary Figures S2b, S3b). As expected, ctDNA fractions were significantly higher in HGSC cases compared to patients with benign tumors (p-values < 0.05, Figure 1A). According to previous findings ^41^, the highest median ctDNA fraction was observed in patients with stage IV disease (p-value = 0.012, Figure 1A). Among patients treated with PDS or NACT, no significant differences in ctDNA fraction were observed (*p* = 0.29, Supplementary Figure S5a), consistent with our previous result ^6^. Furthermore, ctDNA fractions did not significantly differ across primary treatment outcomes (*p* = 0.32, Supplementary Figure S5b).

**Figure 1:**
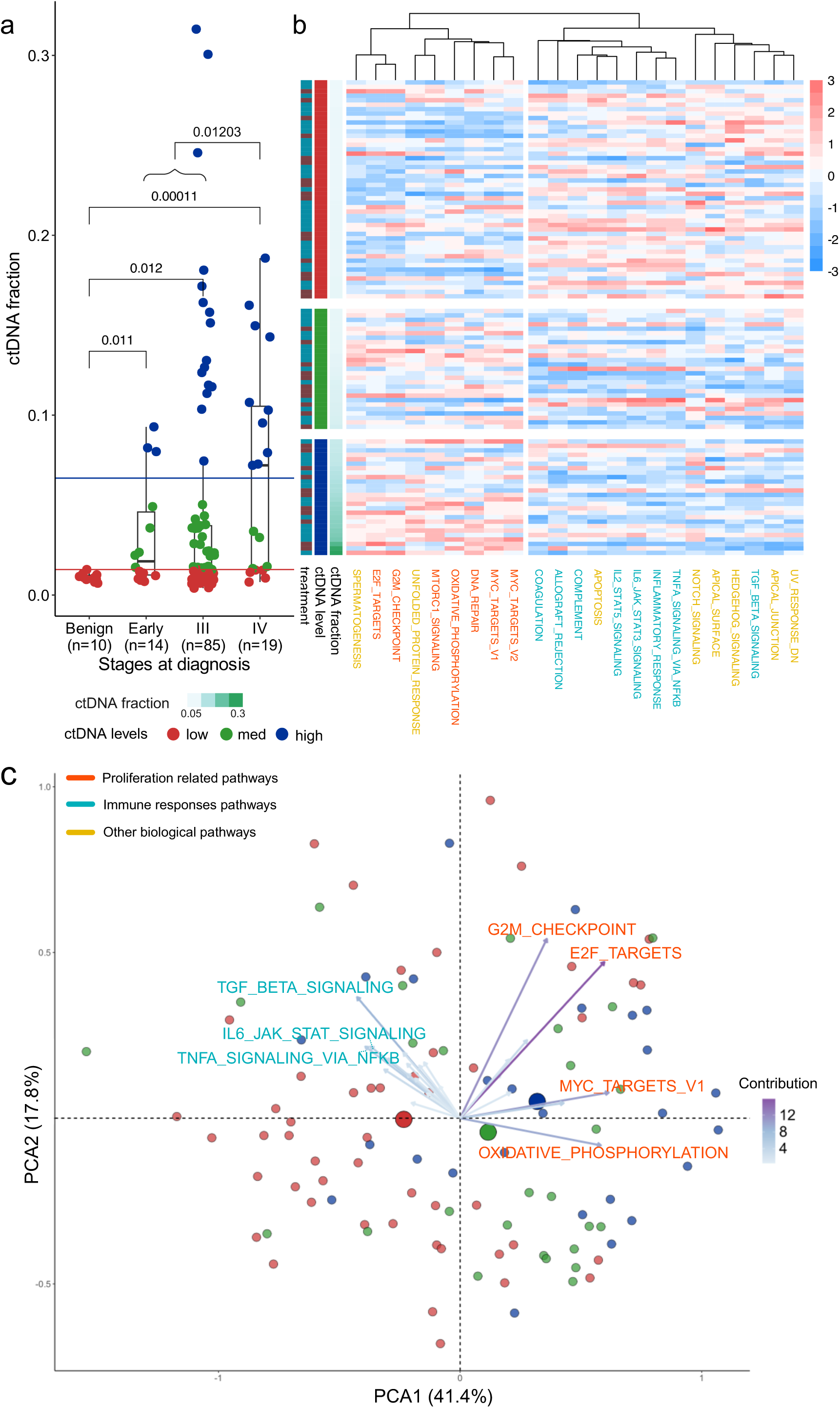
ctDNA level groups have varying expression profiles in tumor tissues. (a) ctDNA levels differ between cancer stages. The red line represents the threshold for low ctDNA, defined by the maximum ctDNA fraction (1.42%) observed in benign (control) samples. The blue line at 6.5% distinguishes high ctDNA from medium ctDNA levels. (b) Expression pathway enrichment differences between ctDNA groups. Heatmap shows enrichment as z-scores, presenting pathways with significant differences between the three groups. (c) Principal component analysis (PCA) of pathway scores, showing the distribution of ctDNA groups based on pathway contributions.

We then stratified patients into ctDNA release groups. The low ctDNA group was defined using the maximum ctDNA fraction observed in benign tumor patients (1.42%) as the threshold (Figure 1A). The high ctDNA level group was defined by ctDNA fractions over 6.5%, consistent with a previous HGSC study by *Paracchini et al.* ^4^. The median ctDNA fraction in our cohort was 1.49%, with 47% of cases classified in the low ctDNA group. No significant differences in clinical characteristics, such as treatment group or age at diagnosis, were observed between the ctDNA groups (Supplementary Table S4).

### Transcriptomic profiles separate low and high ctDNA levels

To investigate the factors influencing ctDNA levels, we first examined the relationship between cancer transcriptomic profiles and ctDNA release. We integrated RNA-seq data from 245 treatment-naïve tumor tissue samples from 102 patients (Supplementary Tables S1, S3) and applied the decomposition method PRISM ^27^ to account for differences in tumor fractions across samples.

First, we performed gene set enrichment analysis (GSEA) using the cancer hallmark pathways from the MsigDB database ^39^. We identified 23 pathways with significant differences between the ctDNA groups (Kruskal-Wallis test, *p-*values < 0.05, Supplementary Table S5). Proliferation-related pathways were more enriched in the high ctDNA group, while immune signaling pathways were more enriched in the low ctDNA group (Figure 1b,c). Particularly, the low ctDNA group was associated with *TGF Beta Signaling* and *TNFA Signaling via NFKB* (PCA contributions ∼8%, Figure 1c).

The medium ctDNA group displayed an intermediate molecular profile between the two extremes. Based on this observation, we next conducted differential gene expression analysis focusing on the difference between the extremes. There were 1,211 significant differently expressed genes (DEGs) between the high and low ctDNA groups (Figure 2a, Supplementary Table S6, *p adj.* < 0.05). Notably, we observed that *CIITA*, required for tumor-intrinsic immunity ^42^, and two membrane-bound *mucins* (*MUC4* and *MUC1*), known for generating cancer-specific immunodominant epitopes ^43^, were expressed higher in the low ctDNA group (Figure 2a, Supplementary Figures S6). Significant differences of these genes were also detected when comparing low group to medium group (Supplementary Tables S7).

**Figure 2:**
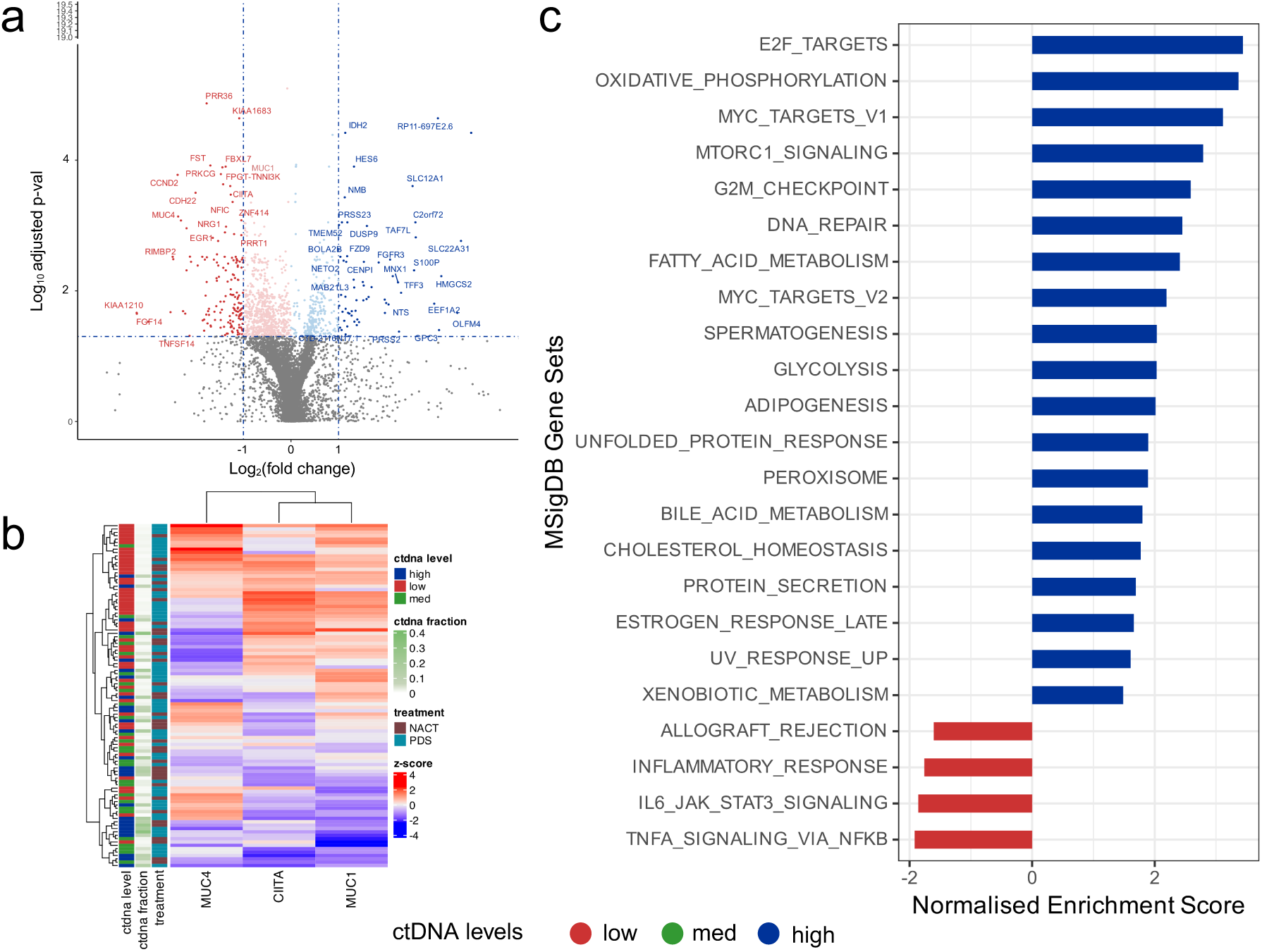
Differential gene expression profiles between low and high ctDNA groups. (a) Volcano plot showing significantly differentially expressed genes between low and high ctDNA release. (b) Heatmap of z-score normalized log2 RNA expression counts for *CIITA*, *MUC1*, and *MUC4*. (c) Pathway enrichment analysis comparing low and high ctDNA release groups. Pathways with normalized enrichment score > 1.5 and adjusted p < 0.05 are shown.

Overall, the contrast between immune and proliferation related pathways between the low and high ctDNA groups was preserved in the two-group comparisons (Figure 2c). Additionally, metabolic pathways including *oxidative phosphorylation*, *fatty acid metabolism*, and *glycolysis* further supported increased proliferative activity in high ctDNA group. The molecular profile of the medium ctDNA group was less distinct (Supplementary Tables S7 and S8), suggesting it may represent a transition state between low and high ctDNA release rather than a biologically separate category.

### Genomic instability correlated with ctDNA levels

Complementary to the transcriptomic profiles, we explored genomic differences in tumor tissues between the ctDNA level groups. We integrated 191 treatment-naïve biopsies whole-genome sequencing data from 100 patients (Supplementary Tables S1, S2). Genomic instability was significantly lower in the low ctDNA group compared to the high ctDNA group, as measured by the number of breakpoints (Figure 3a, Wilcoxon rank test, *p* = 0.038) and mutational burden (Figure 3b, Wilcoxon rank test, *p* = 0.019). To confirm the association between ctDNA levels and both mutational burden and breakpoints, we performed an ANOVA analysis, adjusting for tumor tissue purity, sequencing platform and coverage, HR status, and patient age (Supplementary Table S9). ctDNA levels remained significantly associated with mutation counts but not with breakpoints.

**Figure 3:**
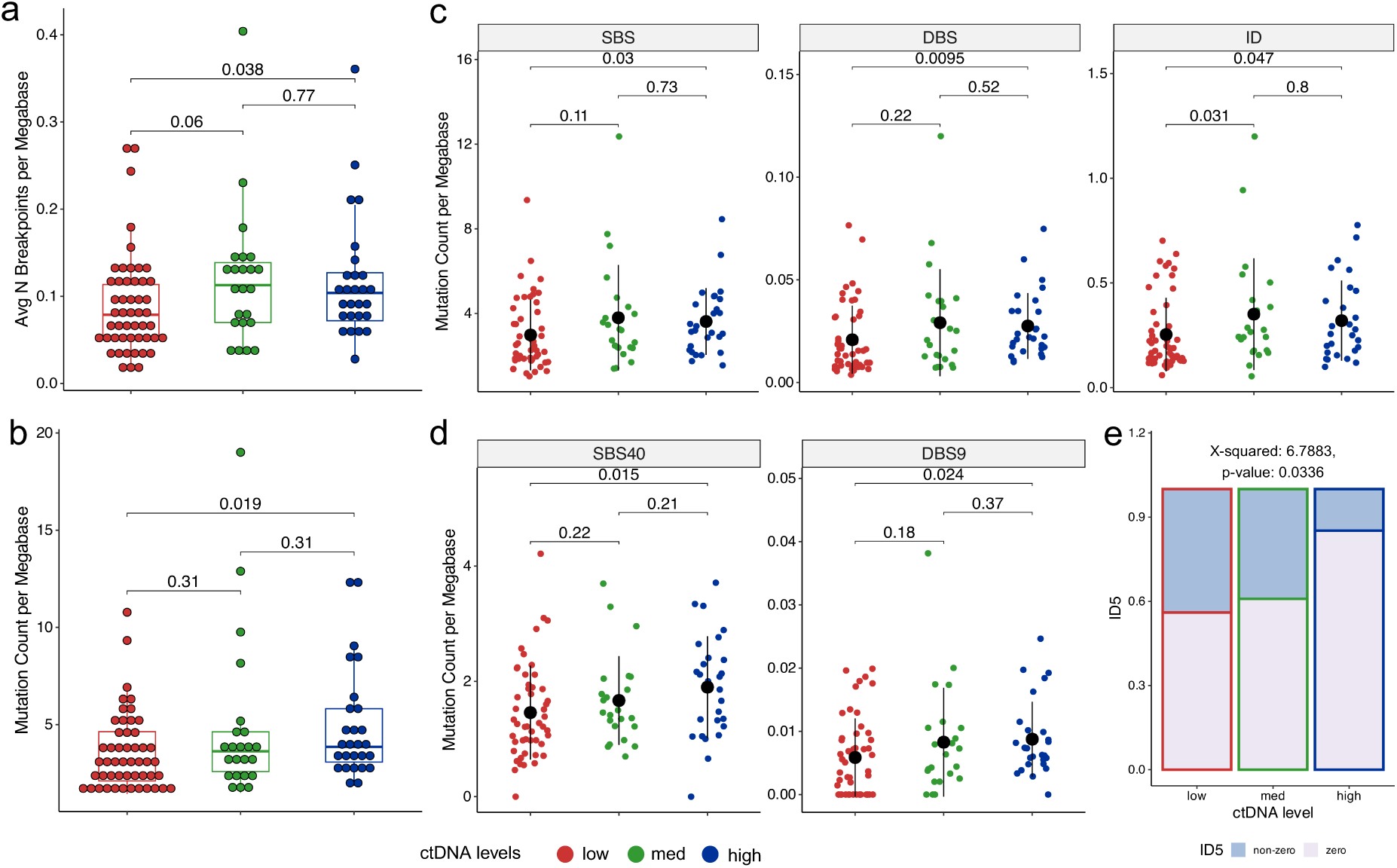
Genomic differences in tumor tissues between ctDNA groups. (a, b) Genomic variant burden compared to ctDNA levels: (a) the number of copy-number breakpoints and (b) the somatic mutational burden per megabase. (c) Mutational burden differences by mutation types: SBS (single base substitutions), DBS (doublet base substitutions), and ID (small insertions and deletions). (d, e) Significant differences between COSMIC mutational signatures by ctDNA levels: (d) mutational burden for two specific signatures, SBS40 and DBS9; (e) the proportion of zero/ non-zero values in signature ID5 across ctDNA levels. See all signature comparisons in Supplementary Figure S9. For pairwise comparisons in (a) and (b), the Wilcoxon rank-sum test was employed. The Kruskal-Wallis test was utilized for analyzing multiple groups. Chi-squared tests assessed the proportion of zero/non-zero ID5 values.

Further analyses of mutation types and mutational signatures (COSMIC) revealed significant differences between the low and high ctDNA groups (Figure 3c,d, *p* > 0.05). The difference in mutational burden could be attributed to two mutational signatures, SBS40 and DBS9, whose mutation counts positively correlated with ctDNA levels (Figure 3d, *p* < 0.05). SBS40 and DBS9 have unknown etiologies, although SBS40 has previously been associated with genomic instability ^44^. In contrast, a negative correlation was observed with the signature contribution of ID5 (Figure 3e, *p* = 0.03), which was more prevalent in patients with low and medium ctDNA levels.

Notably, HR statuses did not differ between the ctDNA groups, specifically, *BRCA1/2* mutations (Supplementary Table S4), or SBS3 (Supplementary Figure S7). No significant group differences were detected between driver gene mutations or copy-numbers (*p*-values > 0.05).

The mutational burden differences highlight the distinct genomic characteristics associated with the extreme ctDNA levels. Similarly to RNA results, medium ctDNA group played as an intermediary phase between the low and high ctDNA groups.

### Prognostic value of ctDNA at baseline differed between treatment modalities

We also evaluated the prognostic value of baseline ctDNA levels across different treatment modalities. Due to recent findings in prognostic differences of ctDNA between NACT and PDS patients ^45^, we analysed these groups separately. ctDNA levels were significantly associated with the risk of progression in NACT patients (Figure 4a, *p* = 0.018) but not in PDS patients (Figure 4b, *p* = 0.55). In PDS patients, cancer stage was main contributor for the prognosis (Figure 4c, Supplementary Figure S8). Surprisingly, in NACT patients, both high and low ctDNA levels were linked to a higher risk of progression compared to patients with medium ctDNA levels at diagnosis (Figure 4d, *p-values* < 0.05). This finding was consistent across both univariate and multivariate Cox regression analyses for NACT patients, with hazard ratios of 4.05 and 5.68 for low and high ctDNA patients, respectively, (Figure 4d, *p*-values < 0.05). Notably, prognostic effect of ctDNA level was independent from the known prognostic value of HRD (Figure 4d*, p=*0.011), defined as in our prior study ^37^.

**Figure 4:**
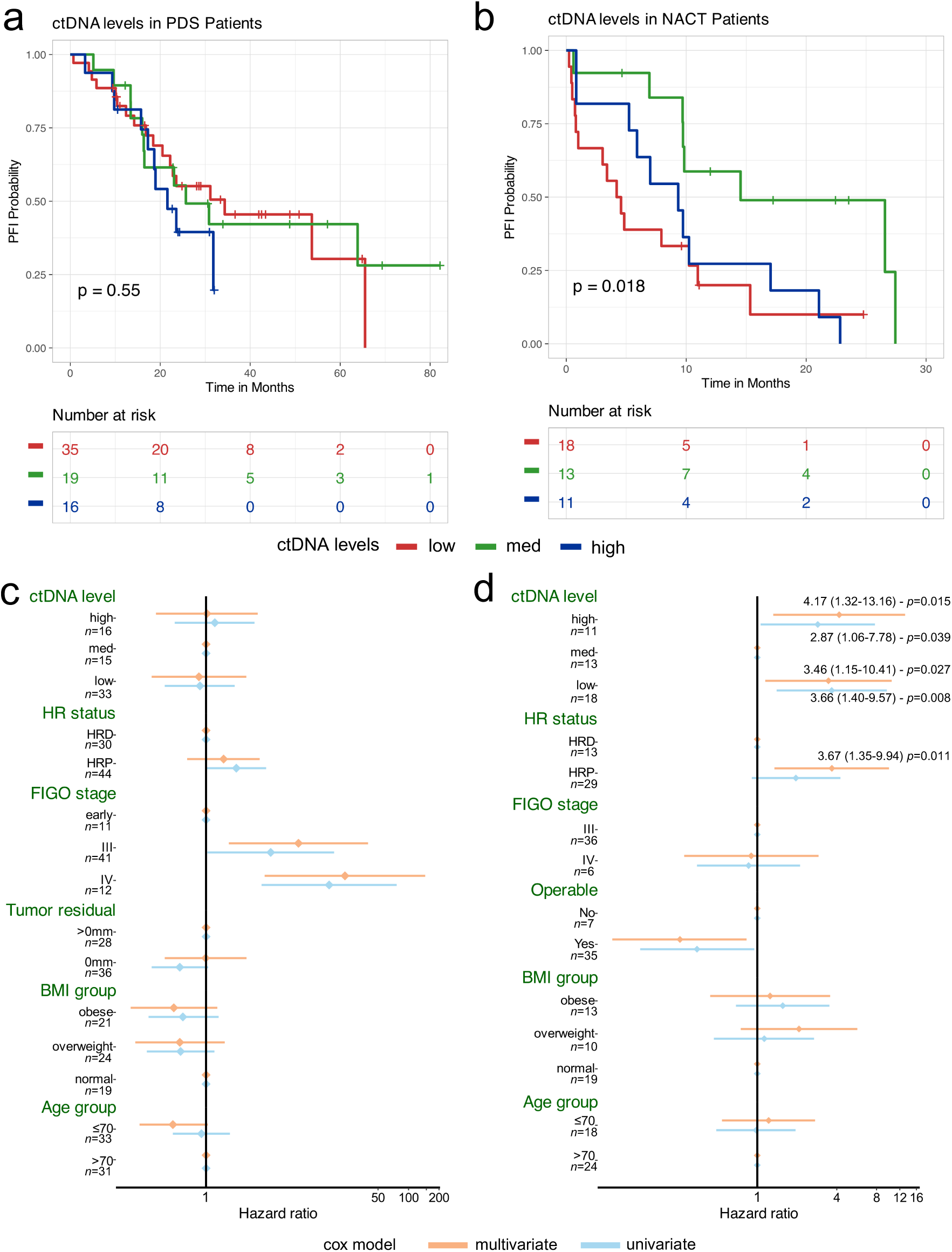
Prognostic value of baseline ctDNA groups. (a, b) Kaplan-Meier survival analysis by treatment type: (a) Patients who underwent primary debulking surgery (PDS) and (b) patients who received neoadjuvant chemotherapy (NACT) before surgery. The log-rank test p-value indicates differences in platinum-free interval (PFI) across all three ctDNA level groups. Five patients (two from the PDS group and three from the NACT group) who did not complete at least two chemotherapy cycles or whose outcomes could not be evaluated were excluded from these analyses. (c, d) Multivariate and univariate Cox regression analysis for (c) PDS and (d) NACT patients, showing associations between PFI and multiple variables, including ctDNA levels. A tumor residual of 0 mm indicates complete resection, while >0 mm indicates incomplete resection. “Operable” NACT patient refers to whether the patient underwent interval debulking surgery after receiving NACT. Homologous recombination (HR) status is categorized as deficiency (HRD) or proficiency (HRP). BMI is classified into three categories: normal (18 < BMI < 25), overweight (25 ≤ BMI < 30), and obese (BMI ≥ 30). Age at diagnosis is grouped into above and below 70 years.

## Discussion

We have characterized the variability in baseline ctDNA levels in patients with high-grade serous carcinoma (HGSC). We identified tumor biology-driven differences in ctDNA levels, mainly related to proliferation and immune responses, and estimated the effect of clinical characteristics. Additionally, we showed that ctDNA level is prognostic only in NACT patients where both low and high ctDNA groups are linked to poor prognosis, which may explain the contradictory findings in prior studies (Table 2).

**Table 2.** Prognostic effect of baseline ctDNA level between different studies.

| Paper | Histology | N (N HGSC) at baseline | Method for detection | Baseline ctDNA amount vs prognosis | Note of the used threshold |
| --- | --- | --- | --- | --- | --- |
| Heo et al. 2024 <sup>45</sup> | Mixed OC | 201 (149) | Targeted sequencing of 9 genes including *TP53* | Baseline differences not reported, poorest prognosis with reduction of ctDNA in detected group. Undetected cases have best prognosis in PDS patients but not in NACT patients. | Threshold 0.2% |
| Azaïs et al., 2025 <sup>10</sup> | Mixed OC | 168 (126) | Targeted sequencing of >500 mutations | No significant differences as univariate. Significant differences when analyzed in BRCA groups (wild type or mutated): undetected ctDNA has shorter OS in both groups. | 0.1% as threshold for undetected ctDNA |
| This study | HGSC | 118 (118) | CNV-based estimation from sWGS | Best prognosis with medium ctDNA with >1.4% and <3% of ctDNA in NACT patients. No differences at PDS patients | Detection threshold 1.4% and high group definition 6.5% |
| Swisher et al. 2005 <sup>57</sup> | Mixed OC | 69 (NA) | Mutant copies per ml from sanger sequencing | Poorer prognosis when ctDNA was detected vs undetected. | Sanger threshold for detection ~5-10% |
| Kallio et al. 2024 <sup>2</sup> | Mixed OC | 56 (44) | Tumor-matched mutation VAF mean from targeted sequencing | Not reported. | Sensitive method, reported detection limit 0.0004% |
| Kim et al. 2019 <sup>33</sup> | HGSC | 41 (41) | Tumor-matched *TP53* mutation from ddPCR | Not reported. | Sensitive method, detection limit ~0.001% |
| Paracchini et al., 2021 <sup>4</sup> | HGSC | 40 (40) | CNV-based estimation from sWGS | Poorer prognosis with high (>6.5%) vs lower ctDNA. | Threshold 6.5% for high, no undetected patients. |
| Rusan et al., 2020 <sup>58</sup> | Mixed OC | 32 (28) | Methylation of *HOXA9* with ddPCR | No significant differences with methylation detected vs undetected. | Sensitive method |
| Pereira et al. 2015 <sup>59</sup> | Mixed gynecologic cancers | 32 (≤22) | Tumor-matched mutations with digital PCR | Not reported. | Sensitive method, detection limit ~0.001% |
| Dobilas et. al, 2023 <sup>60</sup> | Mixed OC | 17 (13) | Tumor-matched mutations with digital PCR | Poorer overall survival in patients with >10 ctDNA mutation copies/ml plasma vs <10 copies. | Sensitive method, detection limit ~0.001% |
| Oikkonen et al. 2019 <sup>3</sup> | HGSC | 12 (12) | Tumor-matched *TP53* mutation from targeted sequencing | No prognostic differences detected at baseline. | Sensitivity <0.5% |
| Parkinson et al., 2016 <sup>61</sup> | HGSC | 7 (7) | *TP53* mutant allele fraction with digital PCR | Not reported. | Sensitive method, detection limit ~0.001% |

OC, ovarian cancer; CNVs, copy-numbers

Tumor biology related factors influencing ctDNA release were studied through transcriptomic analysis of tumor tissues. Low ctDNA was linked to immune related pathways, while high ctDNA level associated with proliferation-related pathways. Accordingly, proliferation-related pathways were identified in previous study of high ctDNA-releasing *HER2*-negative breast cancer patients ^22^. Common pathways such as the enrichment of *E2F targets*, *G2M checkpoint*, *MYC targets* and *oxidative phosphorylation* suggest that increased proliferation relates to high ctDNA amount in both cancers. Additionally, genomic analyses showed that patients with high levels of ctDNA exhibited higher genomic instability, with high contribution of SBS40 signature related mutations. The high mutagenesis profile could be explained by the high proliferation rate, which has been suggested by studies in lung cancer ^20,46^, different breast cancer subtypes ^47^, colorectal cancer ^21^ and neuroendocrine neoplasm ^48^.

Proliferation is known to link to both active secretion of DNA as well as higher cancer cell death ^14^, both possible contributors for enhanced ctDNA release into bloodstream. High proliferation rates are known to induce apoptosis, which explains why ctDNA reflects not only dying cancer cells but also viable tumor populations ^13^. Alternatively, proliferating tumor cells actively release ctDNA into extracellular space via exosome secretion ^14^. Overall, high proliferating tumors have highest ctDNA level in blood, suggesting proliferation as primary source of ctDNA release into bloodstream in HGSC.

In contrast, the low ctDNA was linked to altered immune response. Notably, the low ctDNA group had most prominent overexpression of the gene *CIITA*, linked to tumor intrinsic immunity ^49^, and mucins, known for their role in stimulating immunological epitopes ^43^ and responses ^50^. Additionally, the enrichment of the *TGF Beta Signaling pathway,* which plays a key role in immune evasion ^51^, coupled with activation of *TNFA Signaling via NFKB* pathway, which provides a survival advantage to cancer cells ^52^, suggest altered immune response or microenvironment. Prior studies have suggested that immune cells affect ctDNA release either through clearance of either dying cells or ctDNA, or through cell death mechanisms like rate of necrosis ^14^. For example, macrophages have shown to increase extracellular ctDNA abundance in leukemia cell line culture ^53^. It remains unclear how the observed tumor-intrinsic immune pathways relate to suggested immune mechanisms altering ctDNA, highlighting the need for further studies.

The prognostic value of baseline ctDNA levels in HGSC have presented contradictory evidence in prior studies (Table 2). In our cohort, we detected elevated risk of progression both in high and low baseline ctDNA levels in unresectable patients receiving NACT. Notably, the prognostic significance of ctDNA levels in NACT patients remained independent of homologous recombination status, a known prognostic factor ^37^. In PDS patients, ctDNA levels showed no prognostic value. Furthermore, an NSCLC meta-analysis indicated that adjuvant chemotherapy had limited efficacy in patients with undetectable ctDNA ^54^. Thus, the differences between patient groups and variable thresholds for ctDNA levels may explain the contradictive results between studies, showing poor prognosis for undetected ctDNA in some ^10^ and for higher ctDNA in other studies ^4,22,55^ (Table 2). Overall, low ctDNA levels detected in advanced cancer cases have remained thus far mostly underexplored and are often left out from the analyses. Notably, undetectable ctDNA patients are often excluded from trials requiring ctDNA positivity ^23,24^, limiting research and treatment options on this high-risk group.

We acknowledge the limitations of our study. The high detection threshold in shallow WGS restricted a more in-depth analysis, particularly for among low ctDNA level cases. However, we were able to lower the prior suggested detection threshold of 3% for ichorCNA ^28^ to 1.4%, which we validated by the matched high-coverage sequencing of truncal *TP53* variants (Supplementary Figure S2,S3). Additionally, our results highlight the significant influence of the tumor microenvironment and suggest a potential effect of crosstalk between proliferation and immune pathways. These findings suggest further investigation using additional data layers, such as spatial resolution techniques or empirical approaches to study the release mechanisms.

In conclusion, our study shows that ctDNA release in HGSC is influenced by a complex interplay of proliferation and immune responses. Beyond the prominent effectors, tumor burden, histology, and stage, our findings further explain individual variation in ctDNA levels among HGSC patients. The observation of enriched immune pathways in patients with lower ctDNA levels despite wide-spread disease highlights factors within the tumor microenvironment that impact ctDNA release. This finding can broaden ctDNA usability as a marker of tumor biology even in patients without detectable ctDNA. For instance, patients with negative ctDNA did not benefit from adjuvant therapy in NSCLC ^56^. Understanding the mechanisms behind ctDNA release holds substantial promise for enhancing ctDNA’s clinical utility.

## Declarations

### Ethics approval and consent to participate

All patients participating in the study gave their informed consent, and the study was approved by the ethics committee of the wellbeing services county of Southwest Finland (VARHA/28314/13.02.02/2023).

### Consent for publication

Not applicable

### Availability of data and materials

Sequencing data used in this study can be obtained from European genome-phenome archive (https://ega-archive.org) under studies EGAS50000000674 for sWGS data for ctDNA samples, and for tissues and ascites from EGAS00001004714 for RNA sequencing and EGAS00001006775 for WGS.

The source code for the sWGS pipeline and analysis in this study has been uploaded to https://github.com/maitnnguyen/ctDNArelease_Omics_Integration/tree/main for peer-review. The WGS data analysis pipeline has been published earlier ^26^ and can be found from https://doi.org/10.5281/zenodo.7852210.

## Authors’ contributions

J.H. was responsible for patient recruitment and provided clinical information. J.H., J.O., and M.T.N.N. were in charge of sample selection. M.T.N.N. conceptualized and designed the study, processed sWGS data, developed the bioinformatics pipeline, analyzed, and interpreted the data, and was the primary author of the manuscript and its supplements. D.A., Y.L., and G.M. processed the WGS and RNA data. Y.L. developed bioinformatics pipelines for mutation calling and mutational signatures analysis. K.L. created the bioinformatics pipeline for copy-number calling and tumor purity estimation from WGS data. S.H. and J.O. supervised the study, interpreted the data and edited the manuscript; All authors participated in reviewing the manuscript.

## Acknowledgements

We extend our gratitude to Dr. Kaisa Huhtinen for assistance in sequencing plasma samples, Dr. Ann-Christin Ostwaldt for reviewing the manuscript, and Dr. Kaizhang Zhang for supporting with single-cell RNA preprocessing. We also gratefully acknowledge the computing resources provided by CSC — IT Center for Science.

## Supplement Materials

### Supplementary Method

– Validation of ctDNA fraction estimated by ichorCNA

### Supplementary Tables

– S1: Information of patients and plasma samples in the study

– S2: Information on tissue WGS samples integrated in the study

– S3: Information on tissue bulk RNA samples integrated in the study

– S4: Information of clinical factors by ctDNA groups

– S5: Information on Hallmark pathways and annotation of biological processes

– S6: List of differently expressed genes between high and low ctDNA groups

– S7: List of differently expressed genes between medium and low ctDNA groups

– S8: List of differently expressed genes between high and medium ctDNA groups

– S9: ANOVA analysis to evaluate effects on the mutation count and number of breakpoints

### Supplementary Figures

– S1: DNA amount variation in plasma samples by sample collection year and by sequencing batch.

– S2: ctDNA fraction correlations between ichorCNA-based and *TP53* VAF-based tumor content estimates for samples with less than 3% tumor using two different panel-of-normals.

– S3: ctDNA fraction correlations between ichorCNA-based and *TP53* VAF-based tumor content estimates for samples with less than 6.5% tumor using two different panel-of-normals.

– S4: Correlation between the cancer cell proportion estimated from bulk RNA and copy-number-based purity from matched WGS samples.

– S5: ctDNA fraction by treatment modalities and outcomes.

– S6: Volcano plot from different gene expression analysis between medium and low ctDNA groups.

– S7: Mutational signatures between ctDNA groups.

– S8: Association between complete tumor resection and cancer stage with risk of progression in patients undergoing primary debulking surgery.

## List of abbreviations

HGSC: High-grade serous carcinoma
cfDNA: cell-free DNA
ctDNA: circulating tumor DNA
WGS: Whole-genome sequencing
sWGS: shallow whole-genome sequencing
CNV: Copy number variations
PDS: Primary debulking surgery
NACT: Neoadjuvant chemotherapy
IDS: Interval debulking surgery
FIGO: International Federation of Gynecology and Obstetrics
EOC: Epithelial ovarian cells
DEG: Differently expressed gene
ORA: Overall representation analysis
HRD: Homologous recombination deficiency
HRP: Homologous recombination proficiency
PoN: Panel of normals
NSCLC: Non-small cell lung cancer

