## Supplementary Figures for "Multi-Omics Characterization of ctDNA Release Mechanisms in Ovarian Cancer"

**Figure S1:** DNA amount variation in plasma samples by sample collection year and by sequencing batch.
**Figure S2:** ctDNA fraction correlations between ichorCNA-based and *TP53* VAF-based tumor content estimates for samples with less than 3% tumor using two different panel-of-normals.
**Figure S3:** ctDNA fraction correlations between ichorCNA-based and *TP53* VAF-based tumor content estimates for samples with less than 6.5% tumor using two different panel-of-normals.
**Figure S4:** Correlation between the cancer cell proportion estimated from bulk RNA and copy-number-based purity from matched WGS samples.
**Figure S5:** Comparison of ctDNA fraction by treatment modalities and outcomes.
**Figure S6:** Volcano plot from different gene expression analysis between medium and low ctDNA groups.
**Figure S7:** Mutational signatures between ctDNA groups.
**Figure S8:** Association between complete tumor resection and cancer stage with risk of progression in patients undergoing primary debulking surgery.

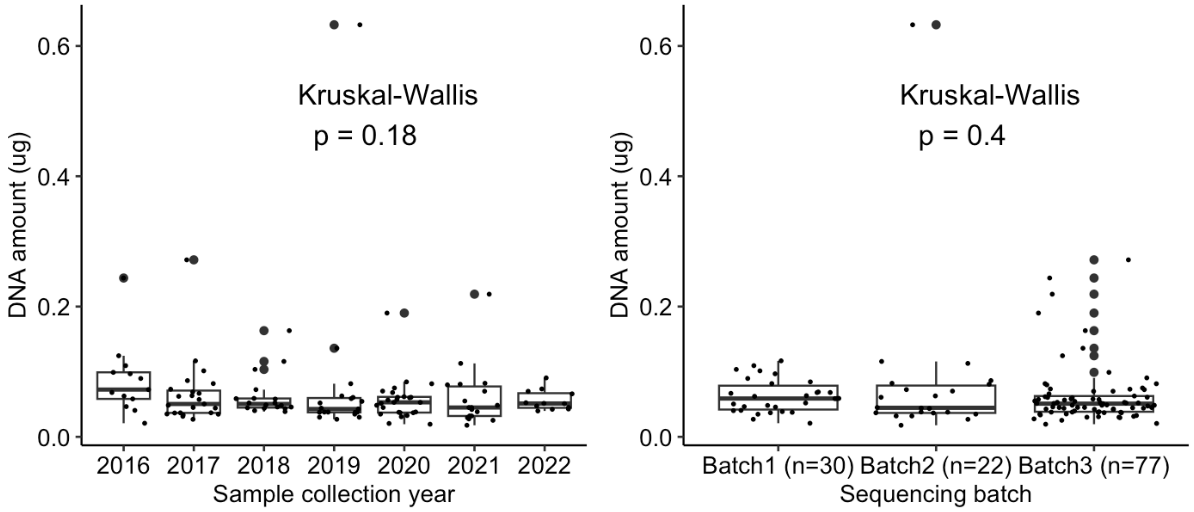


**Figure S1:** DNA amount variation in plasma samples by sample collection year and by sequencing batch.

Plasma samples used in this study were collected between 2016 and 2022 and sequenced during the period of 2022–2023. Here, we evaluated whether DNA amount was affected by the time samples were collected or by sequencing batch. No significant differences related to either sample age or the year of collection/sequencing were found. This ensured the comparability of the plasma samples.

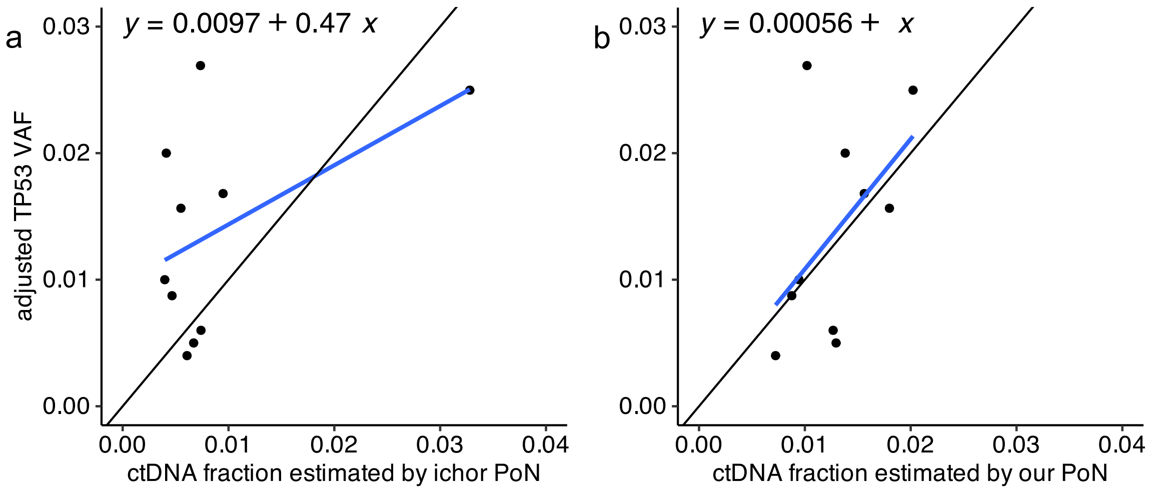


**Figure S2:** ctDNA fraction correlations between ichorCNA-based and *TP53* VAF-based tumor content estimates for samples with less than 3% tumor using two different panel-of-normals. *TP53* VAF values were adjusted for copy-number detected at *TP53* in tissue samples to estimate tumor content more reliably. All samples with adjusted *TP53* VAF ≥ 0.03 were included in the figure. (a) Correlation using PoN reference from the ichorCNA GitHub repository. (b) Correlation using custom PoN constructed from our own white blood control samples. The latter (b) was used to produce the data in this study. Black line represents perfect alignment of estimates and blue line linear regression equation from the data.

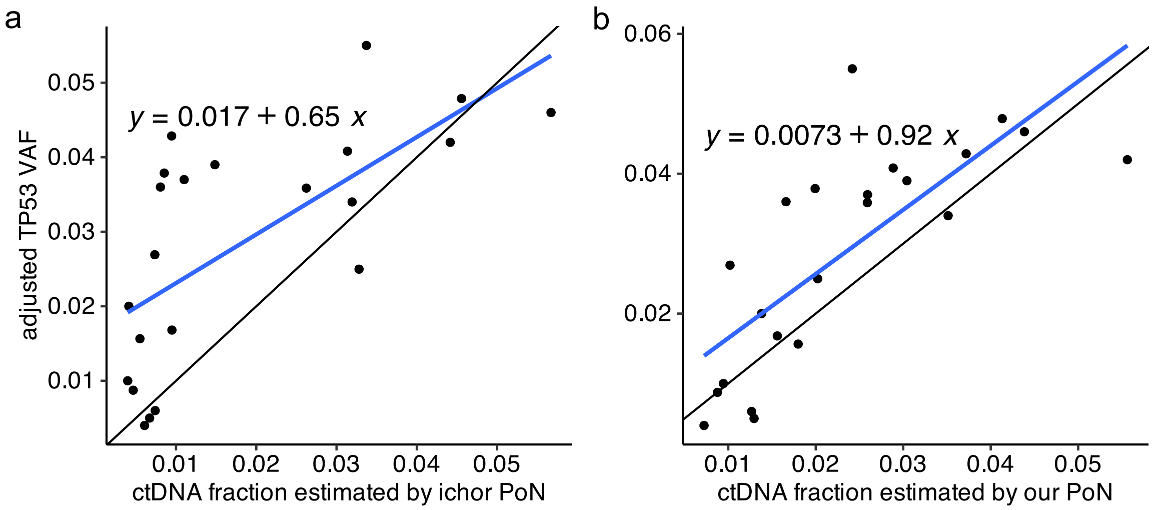


**<u>Figure S3:</u>** ctDNA fraction correlations between ichorCNA-based and *TP53* VAF-based tumor content estimates for samples with less than 6.5% tumor using two different panel-of-normals. *TP53* VAF values were adjusted for copy-number detected at *TP53* in tissue samples to estimate tumor content more reliably. All samples with adjusted *TP53* VAF $\geq$ 0.065 were included in the figure. Linear regression is shown for (a) the PoN reference from ichorCNA GitHub repository, and (b) the custom PoN constructed by our own white blood control samples. The latter (b) was used to produce the data in this study. Black line represents perfect alignment of estimates and blue line linear regression equation from the data.

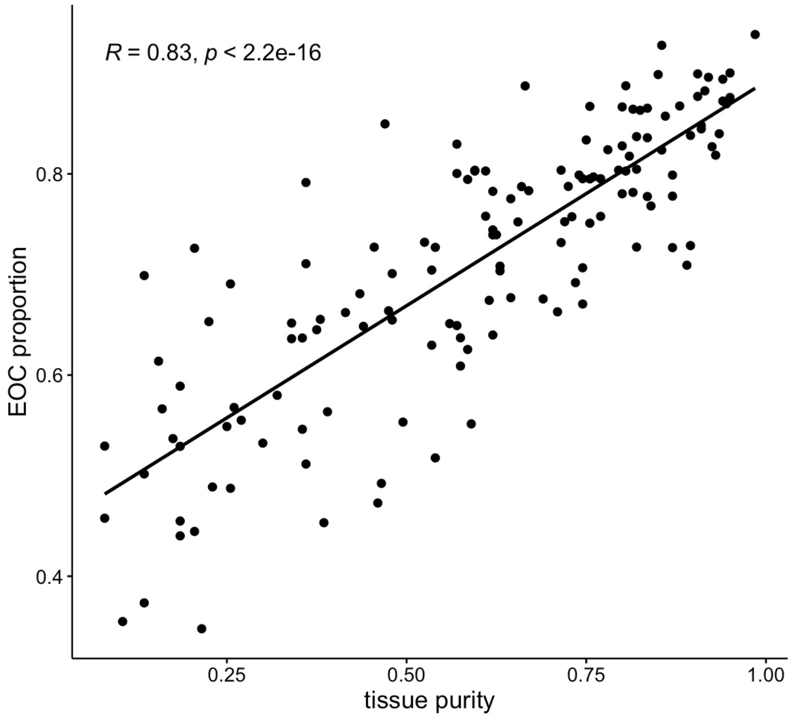


**<u>Figure S4:</u>** Correlation between the cancer cell proportion estimated from bulk RNA and copy-number-based purity from matched WGS samples. EOC: epithelial ovarian cancer

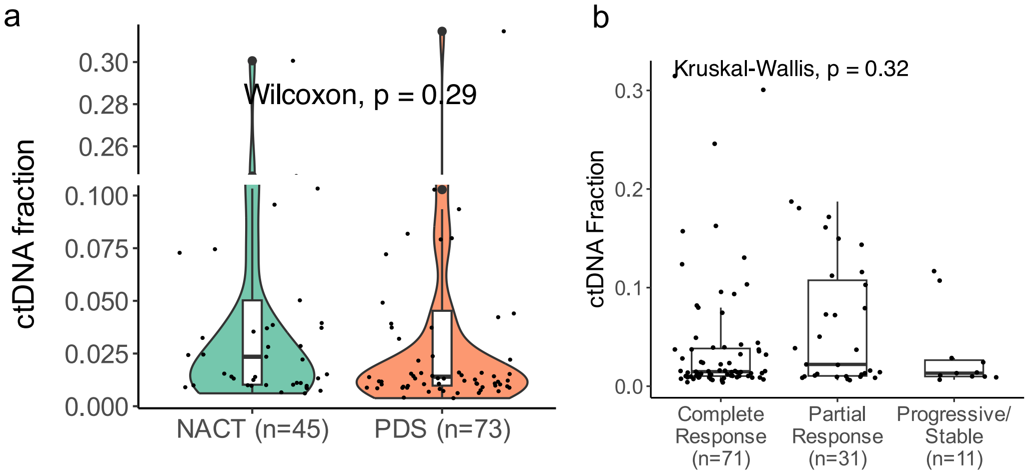


**<u>Figure S5:</u>** Comparison ctDNA fraction by (a) treatment modalities and (b) outcomes

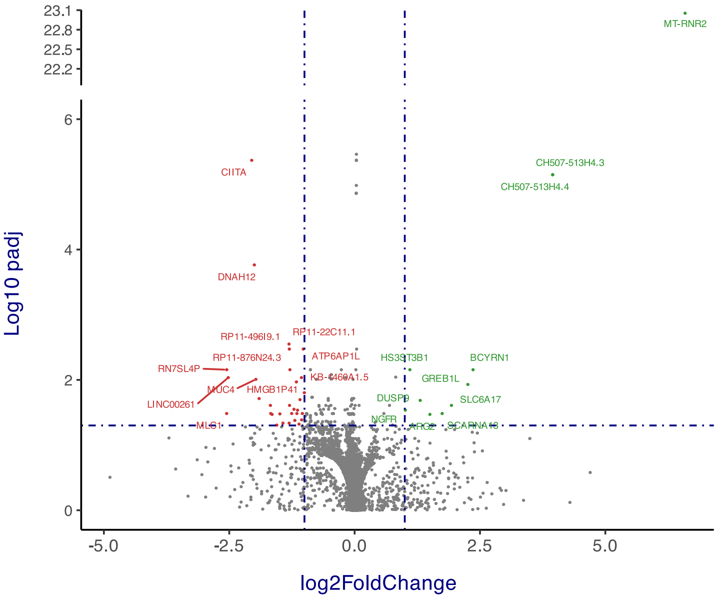


**<u>Figure S6:</u>** Volcano plot from different gene expression analysis between medium and low ctDNA groups

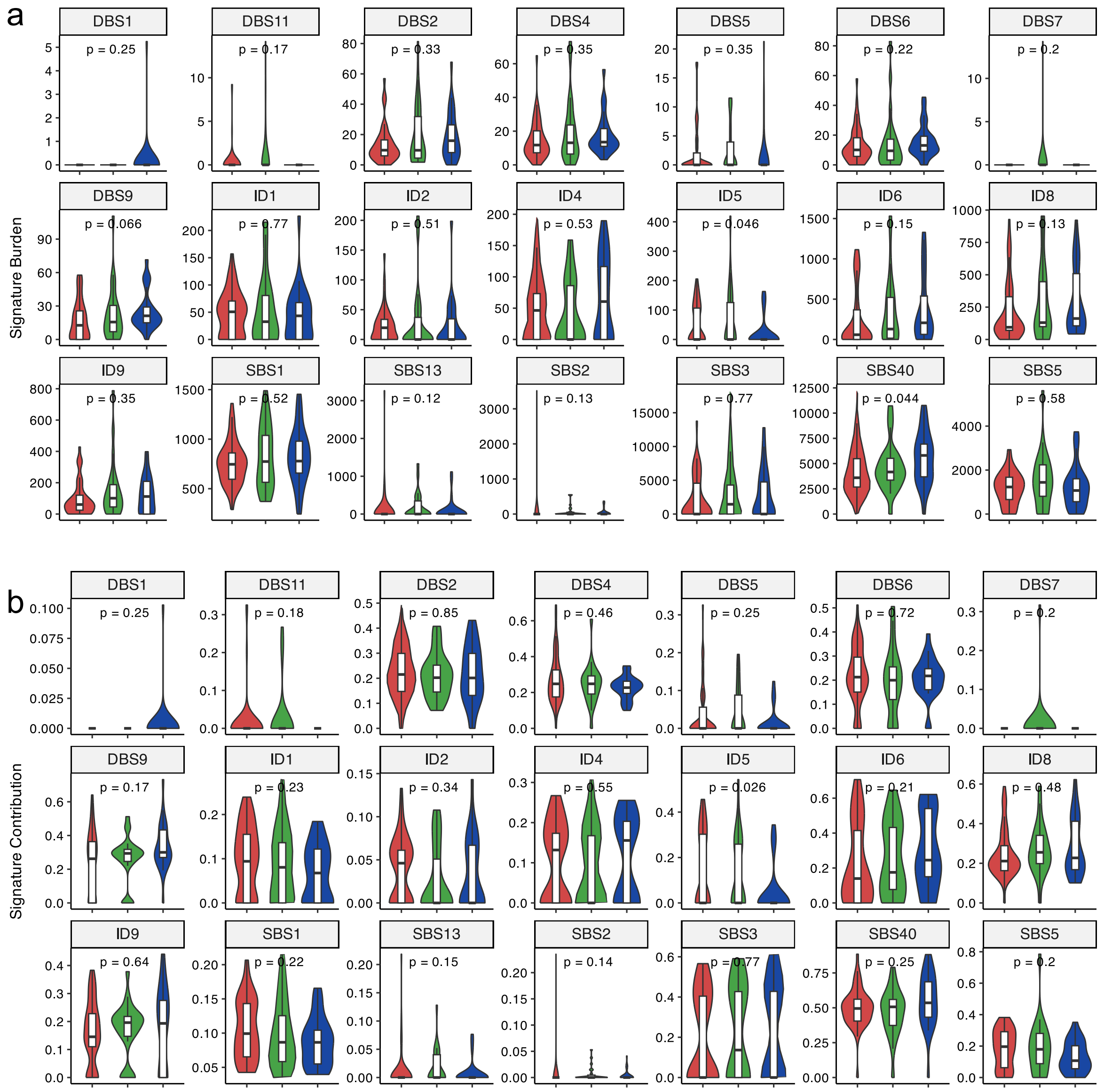


**<u>Figure S7:</u>** Mutational signatures between ctDNA levels.

(a) Comparisons of mutational signature count values by ctDNA levels.

(b) Comparisons of mutational signature contribution values by ctDNA levels.

*Red: low ctDNA group; Green: medium ctDNA group; Blue: High ctDNA group*

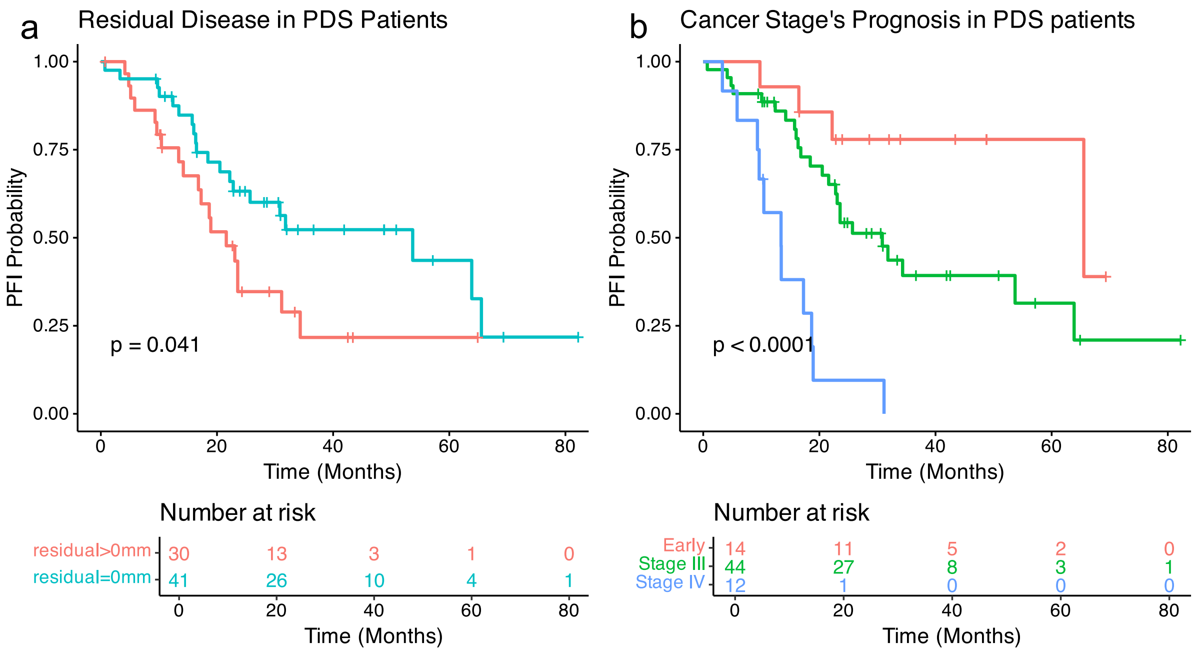


**<u>Figure S8:</u>** Association between complete tumor resection and cancer stage with risk of progression in patients undergoing primary debulking surgery (PDS).
(a) Resection tumor residual.
(b) Cancer stages at diagnosis, Early includes stage I and II.
