## Supplementary Methods for "Multi-Omics Characterization of ctDNA Release Mechanisms in Ovarian Cancer"

I. Validation of ctDNA fraction estimated by ichorCNA

II. Single-cell RNA data analysis

I. **Validation of ctDNA fraction estimated by ichorCNA**

To evaluate the reliability of our estimated ctDNA levels for plasma samples processed with shallow whole-genome sequencing (sWGS), we used truncal *TP53* allele frequency (*TP53* VAF) calling from same plasma samples sequenced additionally with whole-exome sequencing (WES). The *TP53*-based values were used as ground truth. The median coverage of the WES samples was 500x.

To enhance the ichorCNA estimation, we established our own Panel of Normal (PoN) from WGS whole blood control samples to serve as input for the algorithm. This PoN was compared to the original PoN provided by ichorCNA for genome version hg38 on their GitHub page (https://github.com/broadinstitute/ichorCNA/blob/master/inst/extdata/HD_ULP_PoN_hg38_1Mb_median_normAutosome_median.rds).

Pathogenic *TP53* mutations are detected in nearly all high-grade serous carcinoma cases and are used as a measure for inferring tumor purity. We used our bioinformatics pipeline, developed using GATK as described in our previous study, to call *TP53* mutations in tissue WGS samples and force call the selected *TP53* mutations from WES plasma samples. The *TP53* mutations were selected through expert manual curation to verify their pathogenicity and truncal status. Finally, the tumor fraction in plasma samples was calculated from *TP53* VAF, adjusted by the median value of copy number at *TP53* from available tissue samples from the corresponding patient.

We focused to validate the ctDNA estimation for low ctDNA level samples, especially samples with ground truth values smaller than 0.03, threshold recommended by ichorCNA’s authors. Supplementary Figures S2 and S3 present the linear regression of ctDNA fractions between ground truth and estimates using our own PoN or the ichorCNA PoN for samples 0-3% and 0-6.5% ctDNA, respectively. The slope of regression is closer to 1 with our own constructed PoN, which suggests better reliability of ctDNA fraction estimation than with ichorCNA PoN. Notably, only a single sample with <3% ctDNA was estimated to have >1% ctDNA using ichorCNA PoN.

Our PoN was constructed from +200 white blood control samples with median of coverage of 0.7x (provided on github).

II. Single-cell RNA data analysis

To investigate further results from bulkRNA analysis, we collected available single-cell RNA (scRNA) data (Supplementary Table Sx). The preprocessing scRNA data was described in [cite from Anna/Giulia ms]. After that we filtered out samples that have less than 1,000 cell count [citing] to ensure the better resolution followed by at least 5% of cancer cells, annotated based on cancer epithelial cell marker. There is one patient who has more than 1 scRNA sample, we selected the sample that have higher number of cell count. The filtering steps are illustrated in the diagram below.

Markers used for evaluating the expression of MHCII are:
